# Ranking and serial thinking: A geometric solution

**DOI:** 10.1101/2023.08.03.551859

**Authors:** Gabriele Di Antonio, Sofia Raglio, Maurizio Mattia

## Abstract

A general mathematical description of the way the brain encodes ordinal knowledge of sequences is still lacking. Coherently with the well-established idea of mixed selectivity in high-dimensional state spaces, we conjectured the existence of a linear solution for serial learning tasks. In this theoretical framework, the neural representation of the items in a sequence are read out as ordered projections along a suited “geometric” mental line learned *via* classical conditioning (delta rule learning). We show that the derived model explains all the behavioral effects observed in humans and other animal species performing the transitive inference task in presence of noisy sensory information and stochastic neural activity. This result is generalized to the case of recurrent neural networks performing motor decision, where the same geometric mental line is learned showing a tight correlation with the motor plan of the responses. Network activity is then eventually modulated according to the symbolic distance of presented item pairs, as observed in associative cortices of nonhuman primates. Serial ordering is thus predicted to emerge as a linear mapping between sensory input and behavioral output, highlighting a possible pivotal role of motor-related associative cortices in the transitive inference task.

## 1 Introduction

Serial thinking is a cognitive function underpinning almost any of our daily actions. Our brain continuously encodes temporal sequences that we can remember, process, and replay, such as the subsequent steps to make a sandwich or the path from home to work. Serial reasoning underlies logical deduction, categorical and hierarchical thinking, which are crucial to many higher-level cognitive functions (episodic memory, algebraic computation, language, and many others). A still widely open challenge is to find a unique framework in which all these different abilities could be explained, and several hypotheses about sequences mental encoding have been proposed [1, 2].

One of these hypotheses is that sequence information in serial learning tasks is represented in the brain as an ordinal knowledge of the items in the list, independently from their timing or their spatial location. The ranking of the elements of a sequence based on the assigned order implies an abstraction of their arrangement together with the simultaneous encoding of the ordinal and item-specific information [3, 4]. Notably, these are the building blocks of the more general framework named ‘compositional computation’ [5, 6].

Task-relevant information, such as ordinal knowledge in serial thinking, is typically encoded by resorting to specific neuronal representations in the high dimensional state spaces of their activity. These representations in the brain are sparse and involves relatively wide populations of neurons [7, 8]. The high degree of dynamical complexity expressed by such neuronal networks can be exploited to solve relatively difficult cognitive tasks by simply resorting to a linear mapping of their collective neuronal states [9, 10, 11].

Within this theoretical framework, we conjecture the existence of a geometrical solution for serial learning tasks associated with the reorganization of a specific neuronal subspace. This manifold is a line where the arbitrary elements of a sequence can be ranked in any order as suited projections of their neural representations. We derive its analytical form hypothesising it is the so-called ‘mental line’ capable to solve serial ordering tasks [12, 13, 14, 15]. The proposed ‘geometric mental line’ (GML) is capable to explain all the behavioral effects observed in Humans and other animals performing an implicit serial learning task, the so-called Transitive Inference (TI) task [16, 17]. We found that the noisy representation of the stimuli to be ordered plays an important role in learning and shaping the behavioral performances. Our GML is embedded in the low-dimensional space determined by the neural states encoding the items in a sequence, and it turns out to be naturally implemented in recurrent neural networks (RNN). In this framework, the dynamical drift along the GML determining the behavioral output, results to be an attractive manifold where single units strongly correlate, further reducing the dimensionality of the latent state space where the neural activity unfolds.

## 2 Results

### 2.1 Ranking abstract items: The geometric mental line

Consider for example *M* = 4 items *S*_1_ = ‘A’, *S*_2_ = ‘B’, *S*_3_ = ‘C’ and *S*_4_ = ‘D’. Arbitrarily ranking this set of items means to put them in sequences like ‘ABCD’ or ‘CADB’, eventually assigning to each *S*_*k*_ the chosen rank *r*_*k*_ ∈ ℤ, with index *k* ∈ [1, *M*]. In the former example we then have 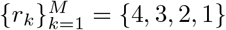, while in the latter the ranks are {3, 1, 4, 2}. A typical example with *M* = 7 often used in cognitive neuroscience is shown in Fig. 1a. Given that, what kind of computational capabilities must a network of neurons be equipped with to solve this task? In other words, what is the machinery needed to encode the generic rank of a set of items?

**Figure 1:**
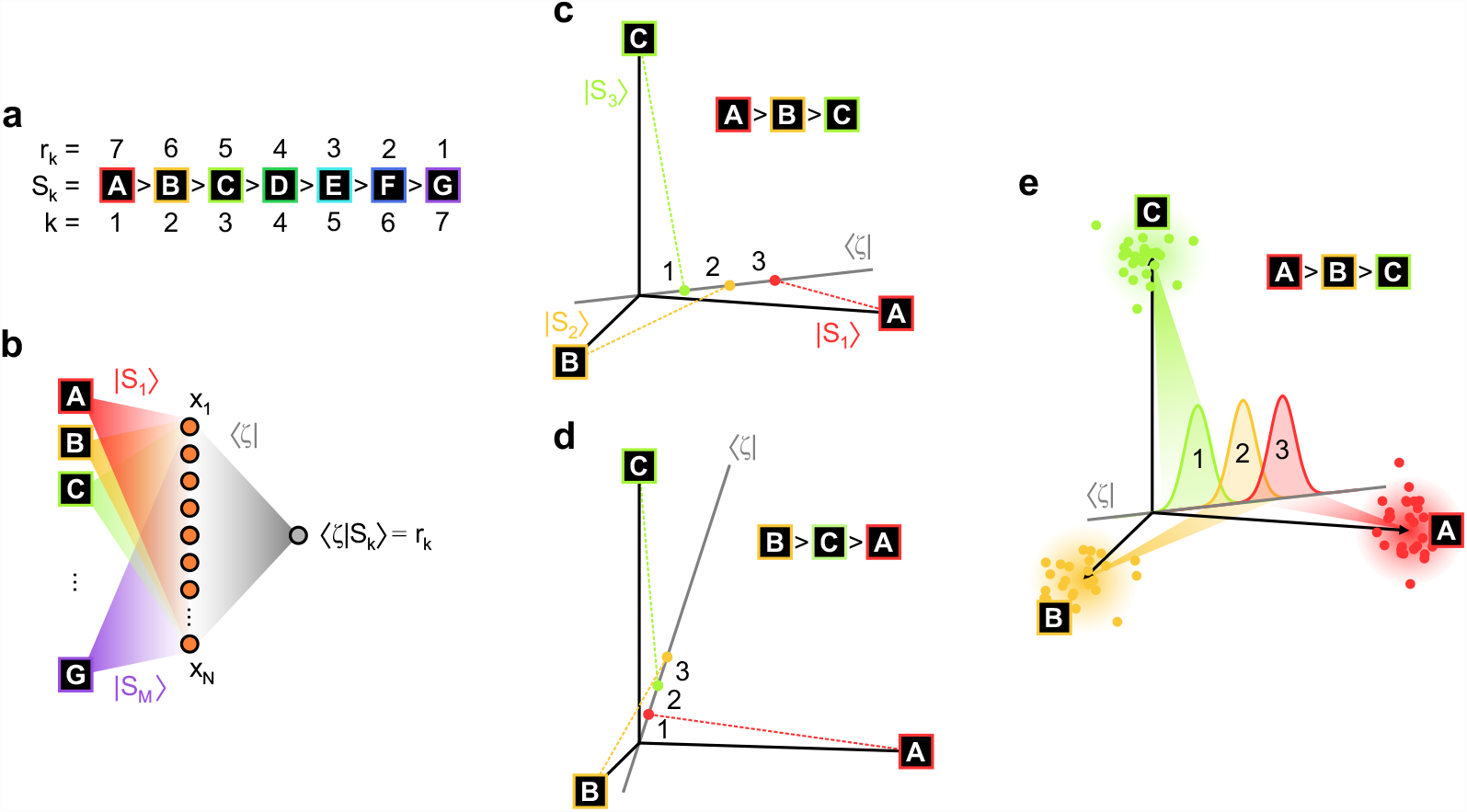
Item representations and the geometric mental line (GML). **a**, An example list of *M* = 7 abstract items *S*_*k*_ here represented as letters (‘A’, ‘B’, …) with rank *r*_*k*_ such that the *k*-th item is greater than *S*_*j*_ only if *r*_*k*_ *> r*_*j*_. **b**, A shallow network (linear Perceptron) with *N* neurons (orange circles) with activity *x*_*i*_ (*i* ∈ [1, *N*]) linearly modulated by the presentation of one of the *M* items in (a). The *k*-isolated item elicits the neural state |*x*⟩ = |*S*_*k*_⟩, which in turn is read out as a projection on the GML ⟨*ζ*| returning the rank *r*_*k*_. Such projection is given by the inner product 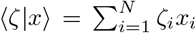 of the Perceptron state and the read-out synaptic weights *ζ*_*i*_ = {⟨*ζ*|}_*i*_ determining the GML. **c**,**d**, Two example sequences with different ordering of three abstract items ‘A’,’B’ and ‘C’ with the related GML ⟨*ζ*| given by Eq. (3). Each item representation lies on the orthogonal axes |*S*_*k*_⟩. Their projections (colored points on the GML) are differently sorted depending on the GML orientation. **e**, Same as (c) but considering a sensory (‘directional’) noise |*ξ*_*n*_⟩ (see main text) in the items representation that now at each trial *n* (colored dotes) are mildly displaced from the corresponding |*S*_*k*_⟩. As a results, projections on the GML will be distributed around the expected rank *r*_*k*_ (colored distributions).

To address this question, we start considering that under stationary conditions, a network of neurons can represent a stimulus (i.e., one of the items *S*_*k*_) as a vector in the space of neuronal states. Referring to the rate-coding framework, this representation corresponds to the set |*S*_*k*_⟩ = |*x*⟩ of firing rates *x*_*i*_ the *N* neurons (*i* ∈ [1, *N*]) have in response to such stimulation (Fig. 1b). For the sake of simplicity here |*S*_*k*_⟩ ∈ ℝ_*N*_ is a column vector where the *i*-th element {|*S*_*k*_⟩} _*i*_ = *S*_*ki*_ can be both positive and negative. As such, neural activity *x*_*i*_ represents the change of firing rate of the *i*-th neuron in a shallow network like a linear Perceptron [18, 19].

In associative cortices, neurons respond to different stimuli with a ‘mixed’ selectivity [20, 21, 7]. Mixed coding favors input decorrelation enabling the flexible storage of relations between items [22], and it is naturally implemented by introducing a layer of randomly connected neurons [23, 7]. According to this, we assume that the inner representations |*S*_*k*_⟩ are independent random vectors such that

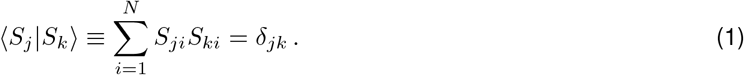

Here, according to the ‘bra-ket’ notation adopted in quantum mechanics [24], ⟨*S*_*j*_| is a row vector given by transposing |*S*_*j*_. ⟨*S*_*j*_ |*S*_*k*_⟩ is the inner product returning the projection of the vector |*S*_*k*_⟩ onto the axis defined by ⟨*S*_*j*_|. This condition naturally holds in the limit of *N* → ∞, provided that *S*_*ki*_ are independent random activities with zero mean and variance 1*/N*. In this limit, Eq. (1) tells us that the set of *M* states |*S*_*k*_⟩ are an orthogonal basis (i.e., the axes) of an *M* -dimensional subspace living in the full *N* -dimensional neural state-space (Fig. 1c).

In this modeling framework, a linear readout unit (the gray circle in Fig. 1b) returning the arbitrary function *f* (*r*_*k*_) of the rank *r*_*k*_ assigned to the *k*-th item,

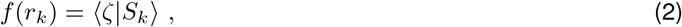

exists and it is given by the following linear combination of items representations:

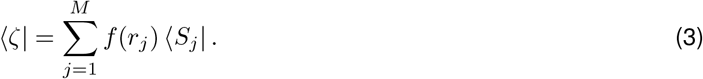

Indeed, from this definition 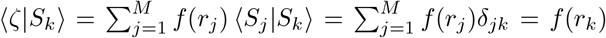 proving Eq. (2) for any set of assigned ranks 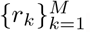. In the particular case *f* (*r*_*k*_) = *r*_*k*_, the row vector of synaptic weights ⟨*ζ*| allows to readout directly the rank of the *k*-th item presented to the shallow network in Fig. 1b. It results to be proportional to the projection of the network state |*x*⟩ along the line oriented as ⟨*ζ*|, as shown in Fig. 1c for the example sequence ‘ABC’. The geometrically composed ⟨*ζ*| is then the searched mental line where items representation are sorted solving a serial ordering task. As such, in what follows we call ⟨*ζ*| the ‘geometric mental line’ (GML). Changing item order from ‘ABC’ to ‘BCA’ as in Fig. 1d, the GML rotates in the *M* = 3-dimensional space of the representations |*S*_*k*_⟩ giving a new set of projections sorted according to the new item position in the sequence.

Now we consider the fact that the neural state encoding an item is unavoidably noisy. This because the activities *x*_*i*_ of units in a network fluctuate due to several sources of ‘thermal’ noise. Indeed, these units aim at modeling cortical cell assemblies composed of a finite number of neurons, each receiving balanced excitatory and inhibitory currents [25, 26]. Another source of stochasticity is the sensory (‘directional’) noise affecting the input received by the network when items are presented. This effect takes into account the fact that item representations are corrupted due to a change of the attentional level, for instance, or a temporary partial view of the visual stimuli. In this case the actual input received by the network can be modeled as |*S*_*k*_⟩ + |*ξ*_*n*_⟩, where the columns vector |*ξ*_*n*_⟩ is a random perturbation occurring at the trial *n*, affecting in turn the direction of the item representation. As shown in Fig. 1e, the result of this sensory noise is that in each trial the item lies in a different position close to its uncorrupted representation |*S*_*k*_⟩. The resulting projections on the GML will be then distributed around the rank *r*_*k*_ with variance 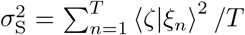 estimated across the *T* trials per item presented. Thus, even under this noisy condition, the GML in Eq. (3) solves on average the serial ordering task.

As we will see later, the stochasticity of both network activity and item representations is a key ingredient to explain the wide set of behavioral effects observed in serial learning tasks.

### 2.2 The Transitive Inference task: learning the GML

Transitive inference (TI) is an implicit serial reasoning task, which consists in generalizing the comparison between items based on an arbitrary assigned rank in a sequence (i.e., A*>*B, B*>*C), that allows to deduct the relationship among stimuli never compared before (i.e., A*>*C). Together with Humans, a plenty of animal species have been found to show such an ability: birds, rodents, fishes, and non-human primates [27, 28]. Successfully performing a TI task thus means having encoded the order of the presented items. Is this ordering computed relying on the same GML derived in Eq. (3)?

To answer this question, we consider as workbench a standard non-verbal TI task, with a list of *M* = 7 items of visual stimuli, conventionally associated with the letters from ‘A’ to ‘G’ and ordered alphabetically as in Fig. 1a. During a first phase of the task, only the learning set of pairs of adjacent items (e.g., ‘AB’, ‘BC’, ‘CD’, …) are presented on a screen in different trials (Fig. 2a). Following the appearance of the pair, the subject is required to choose the item with the highest rank (Fig. 2b). If the item chosen after a reaction time is correct, a reward is received.

**Figure 2:**
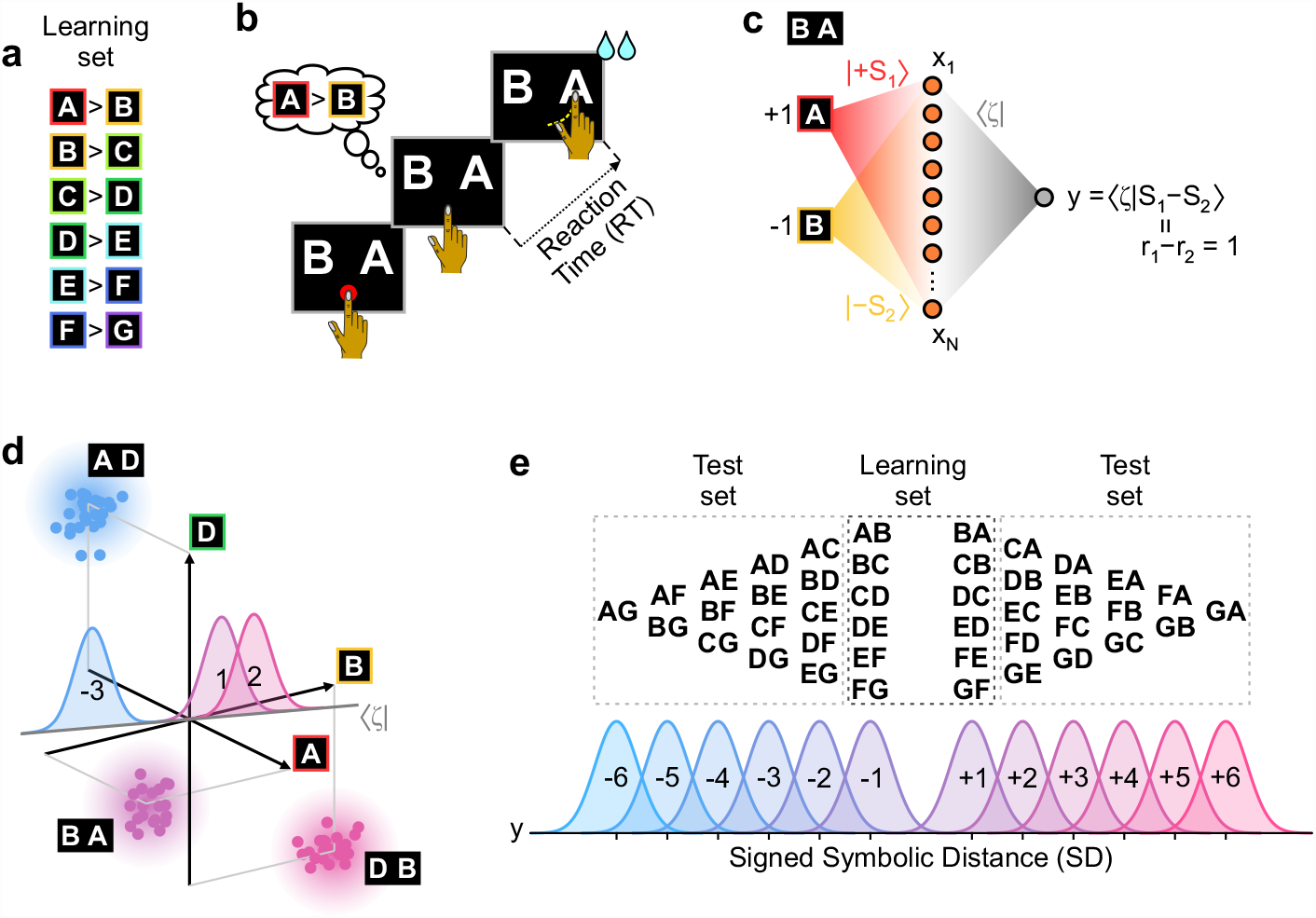
The GML solves the transitive inference task. **a**, The pairs of adjacent items (with their relationship) presented on a screen during the first learning phase of the task. **b**, Example trial of transitive inference (TI) task. Once a pair (BA) is presented on the screen, after the Go signal (disappearance of the red cue) the subject chose to move to the right touching the item with the highest rank (i.e., A). If the choice is correct as in this case, the subject eventually receive a reward. **c**, Shallow network as in Fig. 1 responding to the presentation of the pair ‘BA’ with a positive readout (*y* = 1) instructs to touch the right item (‘A’ with the highest rank as *r*_1_ = 7 *> r*_2_ = 6). The representation |*S*_*k*_⟩ of the presented items (*k* ∈ {1, 2}) are linearly combined as input to the network unit with arbitrary +1 and −1 for the right and the left position, respectively. Readout unit *y* is given by the projection of the network state onto the GML ⟨*ζ*|. **d**, In presence of corrupted items the state vectors |*S*_*j*_ −*S*_*k*_⟩ appear as a distribution of perturbed state (dots). Once projected onto the GML they give raise to Gaussian-distributed readouts centered around the expected signed symbolic distances (SD), which for the example trials ‘BA’, ‘DB’ and ‘AD’ shown are 1, 2 and −3. **e**, Pairs of items presented during the test phase (top) and the expected distributions of readout activity *y* of the shallow network in (c) (bottom). The network ‘response’ *y* is given by the projection of the GML worked out relying only on the item pairs from the learning set (a).

The shallow network in Fig. 2c implements this task by linearly combining the inner representations |*S*_*k*_⟩ of the pair of items appearing on the screen. Coefficients of the combination are arbitrarily set to +1 and −1 to account for the right and left location of the items, respectively, leading to the network state |*x*⟩ = |*S*_*k*_⟩− |*S*_*j*_⟩ ≡ |*S*_*k*_ − *S*_*j*_⟩. The readout unit *y* = ⟨*ζ*| *x*⟩ will inform to reach the right or the left item by responding +1 or −1, respectively. We now ask whether a vector weights in the *M* -dimensional subspace of the item representations, 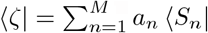 with suited real *a*_*n*_, can solve the task. The set of coefficients *a*_*n*_ should be found taking into account only the learning set of pairs, where the ranks of the presented items always differ by one: |*r*_*k*_ − *r*_*j*_| = 1. Recalling now Eq. (1), the response to a generic pair results to be 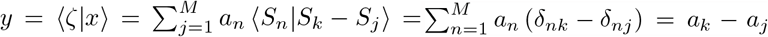, leading to request that for any arbitrary coefficient *a*_*n*_ holds the relationship *a*_*k*_ − *a*_*j*_ = *r*_*k*_ − *r*_*j*_ ∈ {+1, −1} for all item pairs in the learning set. This is a linear system of *M* − 1 equations with *M* unknown variables *a*_*k*_ having an infinite number of solutions *a*_*k*_ = *r*_*k*_ + *φ*, with *φ* any arbitrary real number. Hence, the shallow network solves the TI task with

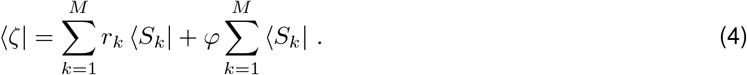

This solution is the same as the GML in Eq. (3) with *f* (*r*_*k*_) = *r*_*k*_ provided that the arbitrary shift proportional to *φ* is removed (i.e., *φ* = 0). Notably, this family of mental lines allows to do the right choice even when the test set of item pairs (unseen during learning) are presented. Indeed

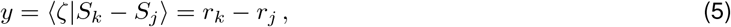

which is the so called ‘symbolic distance’ (SD = *r*_*k*_ − *r*_*j*_) giving for any pair {*S*_*k*_, *S*_*j*_} the response to move to the rightmost item if *y >* 0, i.e., if *r*_*k*_ *> r*_*j*_. Of course if *y <* 0 the item to choose is the one on the left. The learning set of pairs then allows in principle to find the GML solving both the TI task and the serial ordering task introduced in the previous Sect. 2.1. Here it is important to remark that the particular map *f* (*r*_*k*_) = *r*_*k*_ is tightly related to the specific learning protocol adopted in the TI task. Indeed, if a different set of item pairs would be presented during learning or the amount of reward would depend on the serial position of the involved items, a *f* (*r*_*k*_) ≠ *r*_*k*_ would result in Eq. (4).

Notably, the same GML allows to readout another relevant quantity in the TI task: the ‘joint rank’ JR = *r*_*k*_ + *r*_*j*_, i.e., the sum of the ranks of the two presented items. Indeed, by summing the item representation the readout unit gives

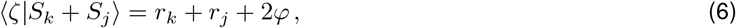

which in the general case of *φ* ≠ 0 leads to have projections on the GML containing a biased information of the joint rank.

Note that in the subspace of the item representations, pairs of items are no more aligned to the |*S*_*k*_⟩ ‘axis’. Indeed, as shown in Fig. 2d, their representation |*S*_*k*_ − *S*_*j*_⟩ bisect the plane determined by |*S*_*k*_⟩ and |*S*_*j*_⟩ of the two presented items. Considering now the sensory noise previously introduced, these representations are randomly distributed as a cloud centered in |*S*_*k*_ − *S*_*j*_⟩. This because in a given trial *n* the actual representation of the pair will be 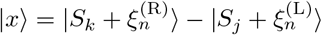. As in Sect. 2.1, the representational noise of left 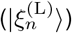 and right 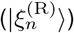 items once projected on the GML are independent random variables with mean 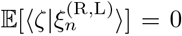 and variance 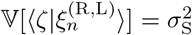. Hence, the readout of noisy pairs is 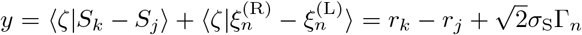, where in each trial *n*, Γ_*n*_ is an independent random Gaussian variable with zero mean and unit variance. One example pair from the learning set (‘BA’ with SD = +1) and two pairs from the test set (‘AD’ and ‘DB’ with SD = −3 and +2, respectively) are shown Fig. 2d, together with the expected Gaussian distribution of the readout symbolic distances.

The distributions of readouts for all the item pairs (i.e., from both learning and test set) are shown in Fig. 2e. In agreement with the so-called ‘symbolic distance effect’ [29], pairs with large |SD| lead our shallow network to respond correctly with higher probability. This because the related Gaussian distribution of the readout *y* is far from the origin. In contrast, the likelihood to make a mistake increases with decreasing symbolic distance, as the chance to have a readout with opposite sign of the rank difference (i.e., (*r*_*k*_ − *r*_*j*_)*y <* 0) widens.

### 2.3 Learning the serial order of noisy items

Although a geometric mental line can be inferred by observing only the pairs of adjacent items in a sequence, and it works on average also in presence of sensory noise, can it be learned in a shallow network of linear units? Modeling effort investigated such possibility proving that this is the case in single-layer networks of formal neurons trained with a ‘delta rule’ [30, 31]. Delta rule learning [32, 33] is equivalent to classical conditioning in behavioral psychology [34, 33] as it reinforces the network weights minimizing the mean square error between expected and actual responses. The learned weights ⟨*ζ*| are then those resulting from a least mean squares algorithm which can be directly computed from the pseudoinverse of the network activity across trials [35].

Following this approach, we computed the learned ⟨*ζ*_L_| in our network performing the TI task in Fig. 2 with noisy pairs of adjacent items, and varying them randomly from trial to trial (see Methods). As expected, the response accuracy (i.e., the fraction of correct responses, Fig. 3a) measured after learning depends on both the difficulty of the task, the level of noise and the rank of involved items (Fig. 3b) [30, 31]. For sufficiently high noise *σ*_S_, performances increase with the symbolic distance (|SD|) between items, giving rise to the so-called ‘symbolic distance effect’ (SDE) in Fig. 3c. Similarly, U-shaped performances clearly arise only for large enough *σ*_S_ displaying higher accuracies for pairs containing terminal items of the sequence Fig. 3d. This is the well-known ‘serial position effect’ (SPE) associated to the fact that first and last items are the ones always winning and losing, respectively, and thus they are always rewarded. From these results a suited noise level 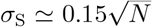 allows to reproduce experimental psychometric functions measured for instance in monkeys [36, 29, 37, 27, 38, 39].

**Figure 3:**
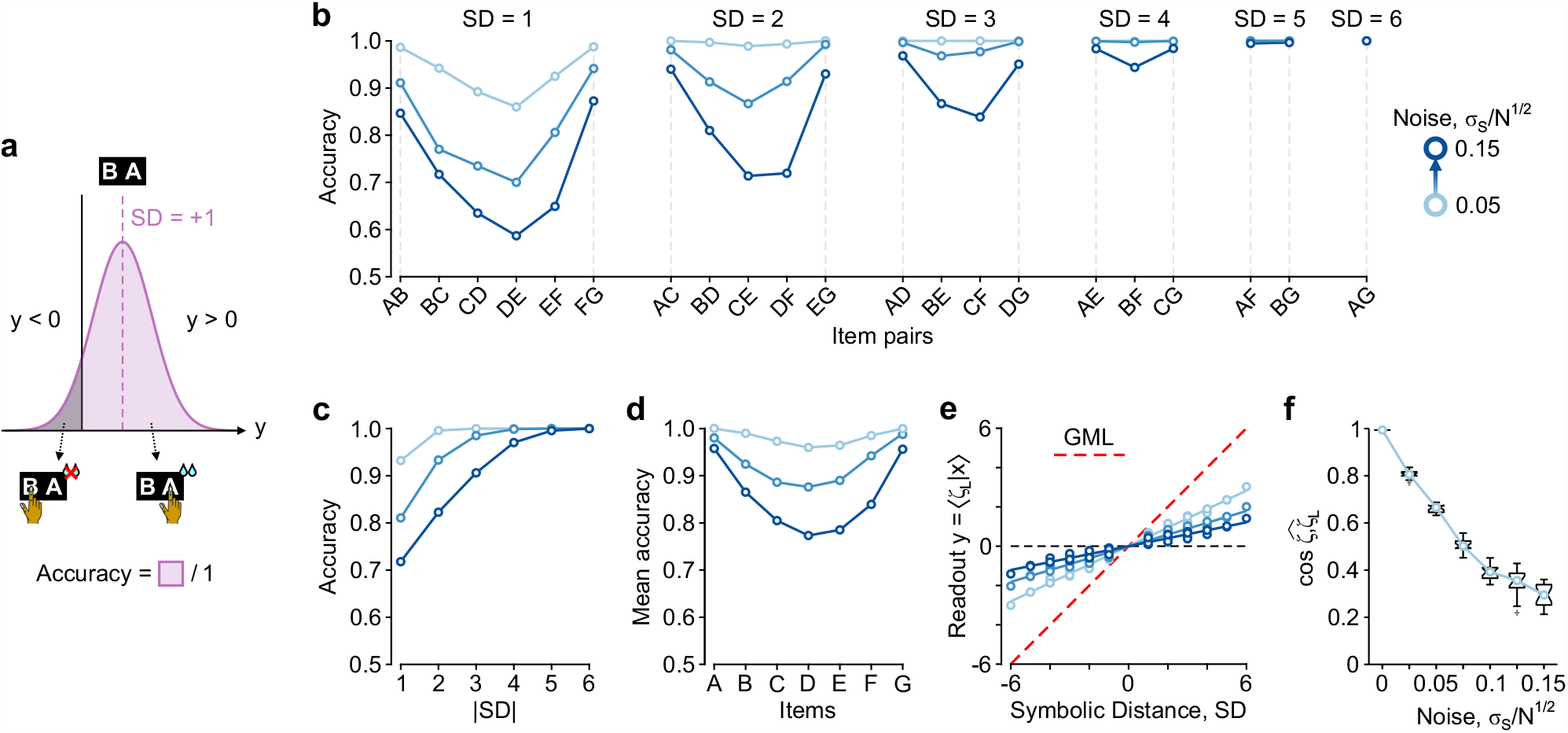
Learning the mental line from the TI task. **a**, Accuracy is the fraction of correct responses *y*, i.e., those pointing to the item with highest rank (SD *y >* 0). **b**, Response accuracy after learning with varying levels of sensory noise *σ*_S_ for item pairs grouped by symbolic distance SD. **c**, Symbolic distance effect (SDE) at varying *σ*_S_. The accuracies in (b) are averaged across pairs grouped by SD in absolute value (i.e., by the easiness of the task |SD|). **d**, Serial position effect (SPE) obtained as in (c) but averaging across items. **e**, Post-learning average readout activity *y* = ⟨*ζ*_L_|*x*⟩ for different SDs and noise levels. Dashed line, *y* resulting from the theoretical GML (*σ*_S_ = 0). **f**, Overlaps between theoretical ⟨*ζ*| and learned ⟨*ζ*_L_| mental lines, i.e., the cosine of the angle 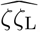. The simulated network (Fig. 2c) has *N* = 1000 units. The *M* = 7 items have sensory noise 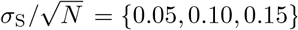. Task sessions have *T/*(*M* − 1) = 25 trials per pairs of adjacent items, and for each noise level *σ*_S_, 20 random sessions are simulated. Weights ⟨*ζ*_L_| for each session are computed from the pseudoinverse of the network activity across the *T* = 150 trials (see Methods). Response accuracies of the learned ⟨*ζ*| are estimated for each item pairs by extracting 10^5^ random realizations of the sensory noise. Box plots in (f) represent the distribution of results across the 20 simulated sessions.

Further inspecting the distribution of responses *y* = ⟨*ζ*_L_ | *x*⟩ across trials with varying difficulty (Fig. 3e), we found that the learned mental line is still a geometric combination of the mean representation of the items. Indeed, a linear relationship between readout activity and symbolic distance is apparent although the slope of the regression now decreases with *σ*_S_. This because the learned GML ⟨*ζ*_L_| is no more parallel to the theoretical one from Eq. (3) (Fig. 3f). In fact the noisy representations |*S*_*k*_ + *ξ*_*n*_⟩ averaged across the presented trials are not fully embedded into the *M* -dimensional space of the exact representations |*S*_*k*_⟩ where the theoretical GML unfolds. Thus, larger is the noise *σ*_S_, smaller is the overlap between these subspaces, meaning that the network state falls in large part into the orthogonal *N* − *M* -dimensional space. As a consequence, the item representation |*S*_*k*_ + *ξ*_*n*_⟩ only in part are embedded in the subspace occupied by the learned GML, leading to *y* values linearly distributed on a range which shrinks with the noise level.

### 2.4 Serial ordering in recurrent neural networks

So far, we solved the transitive inference task relying on a simple one-layer pool of uncoupled units whose activity is readout by a linear Perceptron with weights given by Eq. (3). However, cortical networks have recurrent synaptic connections and the firing rate *x*_*i*_(*t*) of their units is intrinsically stochastic. In this more realistic framework, can the same geometric mental line allow to compute the arbitrary rank assigned to the items of a sequence? To answer this question we consider a recurrent neural network (RNN) with units coupled *via* a random connectivity matrix and each receiving a stochastic input current intended to model an endogenous source of ‘thermal’ noise (see Methods). As for the shallow network in Fig. 2c, the RNN receives as additional input the representations of the items simultaneously presented in pairs (Fig. 4a). Even in this network model, the activity |*x*⟩ is readout as a projection onto the GML (*y* = ⟨*ζ*|*x*⟩).

**Figure 4:**
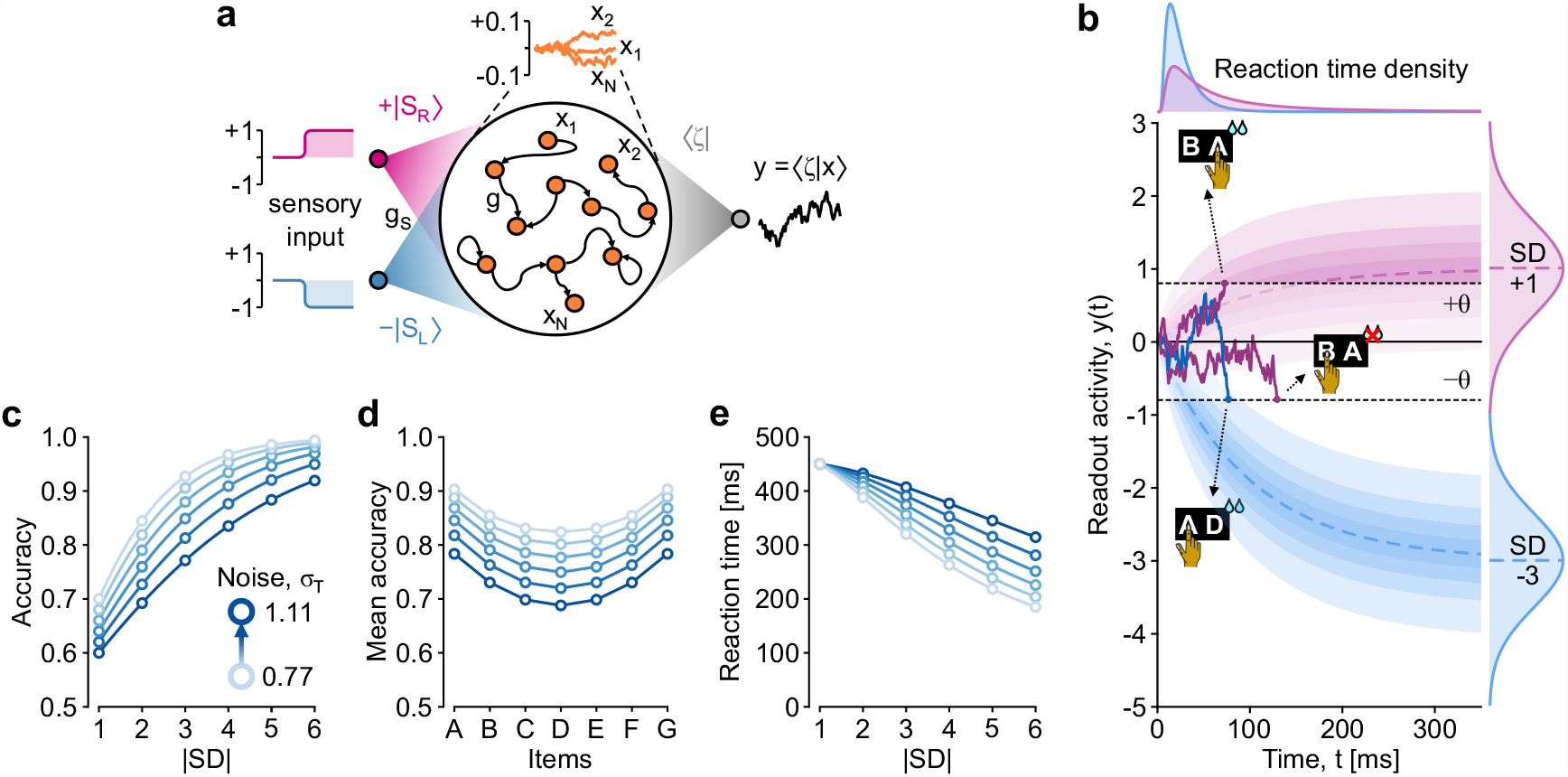
TI task in a recurrent neural network (RNN). **a**, Configuration of a RNN composed of *N* units with stochastic activity *x*_*i*_ (*i* ∈ [*N*]) and random connectivity matrix, modeling a cortical network capable to solve the TI task. Weight of synapses between pairs of units are randomly sampled from a Gaussian distribution with zero mean and standard deviation 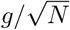. Coupling strength is assumed to be weak (*g* ≪ 1) leading to a linearized dynamics of *x*_*i*_(*t*). As in Fig. 2c, the pair of items presented during a trial provides an input with strength *g*_S_ given by the difference *g*_S_ |*S*_R_ − *S*_L_⟩ between right and left item representations, respectively. The activity randomness is driven by a ‘thermal’ white noise received by network unit as an independent input with same fluctuation size. As in the example shallow networks, the activity |*x*(*t*)⟩ is readout by the unit *y* = ⟨*ζ* |*x*⟩. The vector of weights ⟨*ζ*| is the GML defined by Eq. (3) with *f* (*r*_*k*_) = *r*_*k*_ rescaled in this case by a factor 1*/g*_S_ (see Methods and main text for further details). **b**, Examples of the fluctuating dynamics of the readout activity *y*(*t*) following Eq. (7). The decision is taken when the readout activity crosses for the first time (i.e., the reaction time) one of the two threshold values ±*θ* (dashed lines). The example pairs are ‘BA’ (purple) and ‘AD’ (blue) with symbolic distance SD = +1 and −3, respectively. For ‘BA’ two example trials are shown corresponding to both a correct (choose right) and a wrong (choose left) response. Top, probability densities (p.d.f.) of reaction times for the two example pairs. Shaded areas, deciles of the p.d.f. of *y*(*t*). Right, asymptotic Gaussian densities of *y*(*t*) expected for the two example symbolic distances. **c**, Expected accuracy *α*(SD) from Eq. (8) of the responses at varying symbolic distance and noise level *σ*_T_. **e**, Accuracy per item from (c) averaged across all item pairs of the test set. **f**, Mean reaction time RT(SD) across symbolic distances and thermal noise, setting the relaxation time scale *τ* to have RT|_|SD|=1_ = 450 ms.

Under the hypothesis of relatively weak recurrent connections, the dynamics of the network state |*x*⟩ can be linearized giving rise to the following stochastic differential equation for the readout unit (see Methods):

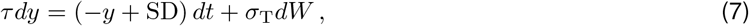

which turns out to be a leaky integrator driven by a Gaussian white noise *dW* (*t*) with zero mean and covariance *dt*, an infinitesimal time step [40]. Here *τ* is the relaxation time scale of the network units and *σ*_T_ is the size of the endogenous thermal noise affecting each unit of the network. Here, the distribution of the readout activity exponentially adapt to a Gaussian distribution centered around the symbolic distance SD between the presented items (Fig. 4b), thus RNN can solve the TI task relying on the same GML derived above.

In this modeling framework, the RNN readout can play the role of the ‘decision value’ in diffusive models of perceptual decision [41]. Accordingly, the decision to reach the right (left) item on the screen is taken when the threshold level +*θ* (−*θ*) is crossed by *y*(*t*) for the first time Fig. 4b [30]. For such a decision process, the accuracy (i.e., the probability to cross the correct decision threshold and thus to make the right choice) and the reaction time (i.e., the time *T* when |*y*(*T*)| *> θ* for the first time starting from *y*(0) = 0) can be analytically derived for *θ* ≪ 1. Indeed, in this limit Eq. (7) is well approximated by a Wiener process with two ‘absorbing’ barriers [40]. For it the probability to have *y*(*t*) crossing the positive threshold *θ* when SD = *r*_R_ − *r*_L_ *>* 0 is given by

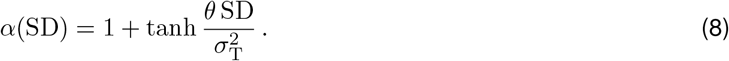

This accuracy is an estimate of the fraction of correct trials where the item on the right is chosen as having the highest rank. Similarly to what found for shallow networks in presence of sensory noise in Fig. 3c, the symbolic distance effect clearly emerges (Fig. 4c). Indeed, the accuracy decreases with task difficulty (smaller |SD|) and size *σ*_T_ of the endogenous noise. The serial position effect is also recovered (Fig. 4d), such that the terminal items in the sequence are associated with the highest accuracies. Despite such similarities, a more careful inspection allows to find a significant difference between the case of sensory and thermal noise. Indeed, performances on terminal items are more affected by *σ*_T_ rather than to *σ*_S_.

The reaction time *T* is the other behavioral output we can workout analytically from the stochastic dynamics (7) of the readout. In this theoretical framework, *T* is the ‘first-passage time’ of the readout activity through the threshold *θ*. It is a stochastic variable whose mean results to be [40]

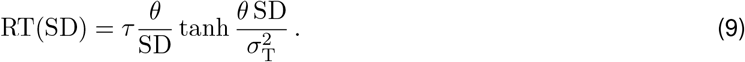

As shown in Fig. 4e, the mean response time reduces with the ease of the decision to take, i.e., with the symbolic distance between the pairs of presented items. This serial position effect is another hallmark of the TI task [30, 42, 27, 39], according to the well-known speed and accuracy trade-off in decision making [41].

### 2.5 Optimal balance between different sources of noise

As sensory and endogenous noise differently affect the behavioral performance of the RNN, we further investigate this aspect in a more realistic scenario. We expose the same linear RNN to a continuous stream of trials where the usual item representations incorporate the sensory noise introduced in Sect. 2.3 (Fig. 5a). In this *in silico* experiment, the network state at the beginning of each trial is no more set to 0, as it is the case in cortical networks. The vector of the readout weights ⟨*ζ*| is learned resorting to the pseudoinverse of the activity matrix aiming at reproducing the right choice in the trials with item pairs from the learning set (*y* = SD = ± 1, see Methods).

**Figure 5:**
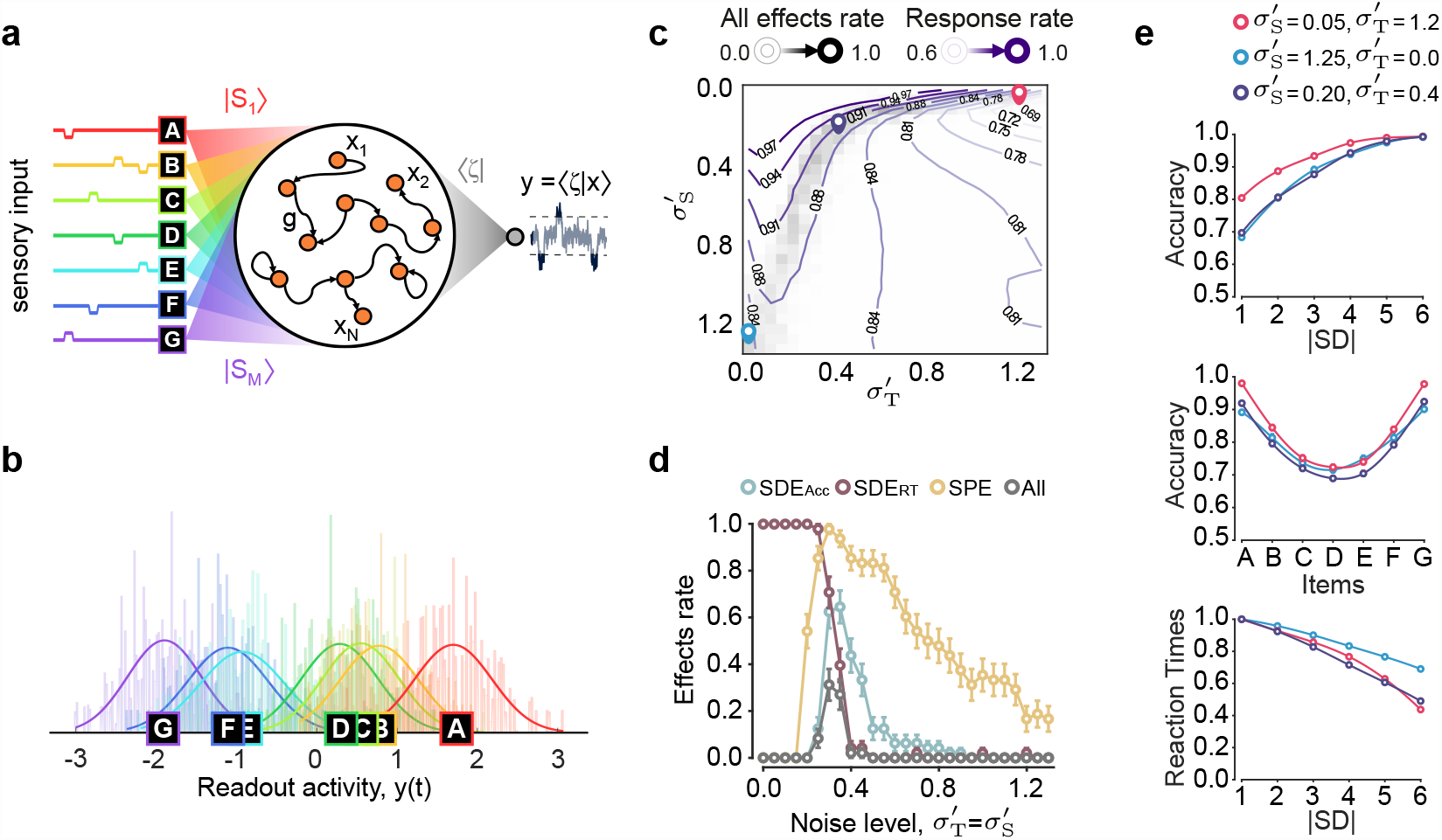
Behavioral effects in TI tasks are reproduced in RNN with balanced noise. **a**, The same linear RNN as in Fig. 4a receiving a continuous stream of randomized trials of the TI task incorporating simultaneously both the sensory and thermal noise (see Methods). **a**, The same linear RNN as in Fig. 4a receiving a continuous stream of randomized trials of the TI task incorporating simultaneously both the sensory (directional, 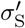) and thermal (endogenous, 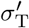) noise. The network is trained on the item pairs of the learning set (|SD| = 1) computing the readout weights ⟨*ζ*| from the pseudoinverse of the network activity assuming a +1 (−1) response when the highest-rank item is on the right (left). When the readout activity crosses for the first time a threshold level (|*y*(*t*)| ≥ *θ* = 0.5) the RNN takes a decision. If *y*(*t*) does not cross such threshold within 1.5 s from the onset of the item pairs, the trial is not taken into account in the computation of the performances. **b**, Distribution of the post-learning readout activity when only one it em is presented per time. **c**, Frequency of network realizations (gray shading) capable to reproduce all the behavioral effects expected for the TI task, by varying the size of both the sources of noise (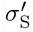 and 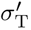). A RNN reproduces such effect if simultaneously displays i) an accuracy and a reaction time increasing with |SD| (SDE_Acc_ and SDE_RT_, respectively) and ii) if the mean accuracy of terminal items is greater than the central ones (SPE). Contours represent the response rate, i.e., the fraction of trials with RNN taking a decision. **d**, Frequency of the behavioral effects (averaged over 50 independent RNN realizations) for varying balanced level of noise 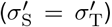. **e**, Behavioral effects (averaged over 50 RNNs) co-occurring in different noise regimes pointed out in (c) by colored cirlces: strongly thermal (red), balanced (cyan) and strongly sensory (purple) noise. The balanced noise case, is the one with the highest frequency of responses.

All these additional sources of uncertainty does not affect the capability of the network to encode the serial order of the items. Indeed, stimulating the trained network with each item presented alone, the readout activity returns separated distribution of values. According to the expectation such values correspond to the projection of the network activity onto our GML, where the items are sorted based on their relative rank (Fig. 5b).

However, inspecting different combinations of noise levels, it is apparent that only a tight balance between them allows to have the simultaneous occurrence of the mentioned behavioral effects (dark gray band in Fig. 5c). This analysis is performed by changing both sensory and thermal noise levels, here denoted by 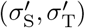. These parameters represent the effect of noise directly on the activity of the network units and they are tightly correlated with the size of noise fluctuations (*σ*_S_, *σ*_T_) measured along the GML (see Methods). The existence of a ‘sweet’ spot for the balance between sensory and endogenous noise, is even more strikingly represented by the narrow peak in the rate of occurrence of all effects along the line 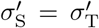 (Fig. 5d). Such balance clearly emerges from the competition of a two-fold role of the noise. A destructive role, occurring when it is so large to disrupt the capability of the RNN to produce the correct response, and a constructive one, as noise is a needed ingredient to have both the symbolic distance and serial position effects (SDE and SPE, respectively).

Although along the narrow band of balanced noise the behavioral effects are similarly reproduced (Fig. 5e), only when both sensory and thermal noise have a comparable size (purple circle, 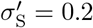 and 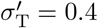) the RNN has the highest rate of responses (i.e., the fraction of trials in which a decision is taken). This is the configuration of balanced noise best fitting experimental evidence, and we use it as reference network for subsequent analyses.

### 2.6 Impact of sequence length and size of the training set

Using the above RNN with balanced noise as an *in silico* experiment, we now investigate the impact on some relevant parameters of the TI task. This to gain further insights and make predictions about how learning of serial ordering can be experimentally modulated.

Firstly we focus on the size of the training set (Fig. 6a). For a small number of learning trials, noise is too high to reach significant performances, limiting the capability to reliably estimate the proper vector of readout weights. This leads to indistinguishable projections of the network activity onto the GML with almost equally low performances and long reaction times (lighter curves). Increasing the number of learning trials, all the behavioral effects are recovered. Interestingly, we found the accuracy displays a change of concavity in the performances and a flattening in the SPE for mid-rank items, which can be exaplained by the sigmoidal dependence highlighted in Eq. (8).

**Figure 6:**
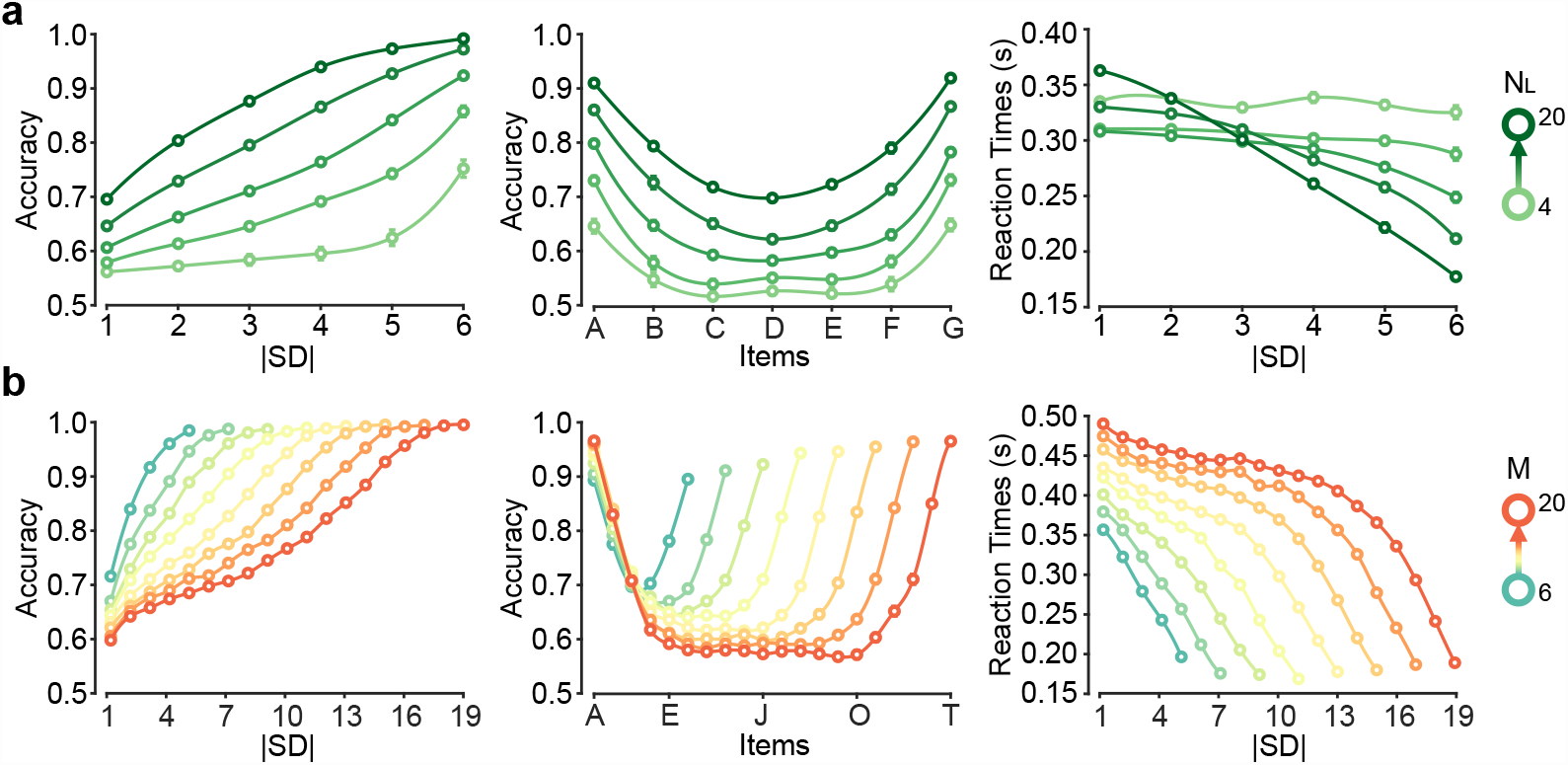
TI task in RNN with varying number of training trials and items to sort. **a**. Behavioral effects: SDE for the accuracy and the reaction time and SPE from left to right, respectively. Number of learning trials range from 4 to 20. Network as in Fig. 5 with 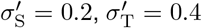. Accuracy and reaction times are averaged across 50 random realizations of the RNN. **b**. Behavioral effects as in (a) for increasing numbers of items in the sequence to sort. Item number range from 6 to 20.

By increasing the number of items to be sorted in the TI task, a strong modulation of the psychometric curves can be also observed (Fig. 6b). SDE displays a change of concavity both in the reaction times and in the accuracyleading to more rapid changes for larger symbolic distances. Another interesting effect is the equalization of the middle-rank representations as soon as the lowest possible accuracy of 0.5 is approached: a barrier limiting the maximum number of items that can be sorted.

### 2.7 Learning the GML in RNN taking motor decisions

So far we studied how the GML ⟨*ζ*| is learned assuming that the only plastic synapses were those between the network units *x*_*i*_ and the readout *y*. Here we investigate a more realistic framework, where the cerebral network involved in a TI task is represented by a RNN encoding the ‘motor’ output by confining its activity |*x*⟩ within a low-dimensional latent space [43]. Such choice follows from the experimental evidence that the premotor cortex (PMC) of monkeys contributes to the motor decision in a TI task with a neuronal activity modulated by the symbolic distance between item pairs [39].

To this purpose, in addition to the sensory input *g*_S_ |*S*_R_ − *S*_L_⟩, the RNN introduced in Fig. 5 receives also the motor-related input *g*_M_ |*μ*⟩ (Fig. 7a) (see Methods). As shown later, this additional input determines the axis along which the network activity is forced to unfold as the decision to move is taken. With this we aim at roughly modeling motor plans known to be encoded in PMC. Here *g*_S_ and *g*_M_ modulate the strength of the sensory and motor-related inputs, respectively. As in previous network configurations, the learning phase involves only the set of pairs {*S*_*j*_, *S*_*k*_} with symbolic distance |SD| = |*r*_*j*_ − *r*_*k*_| = 1 (Fig. 2a), and we impose *y* = +1 (−1) when the item to reach is on the right (left). In this phase the mental line ⟨*ζ*| is computed resorting to the pseudo-inverse of the RNN activity in absence of feedback (i.e., without re-injecting *y* = ⟨*ζ* |*x*⟩ as a factor of |*μ*⟩, see Methods).

**Figure 7:**
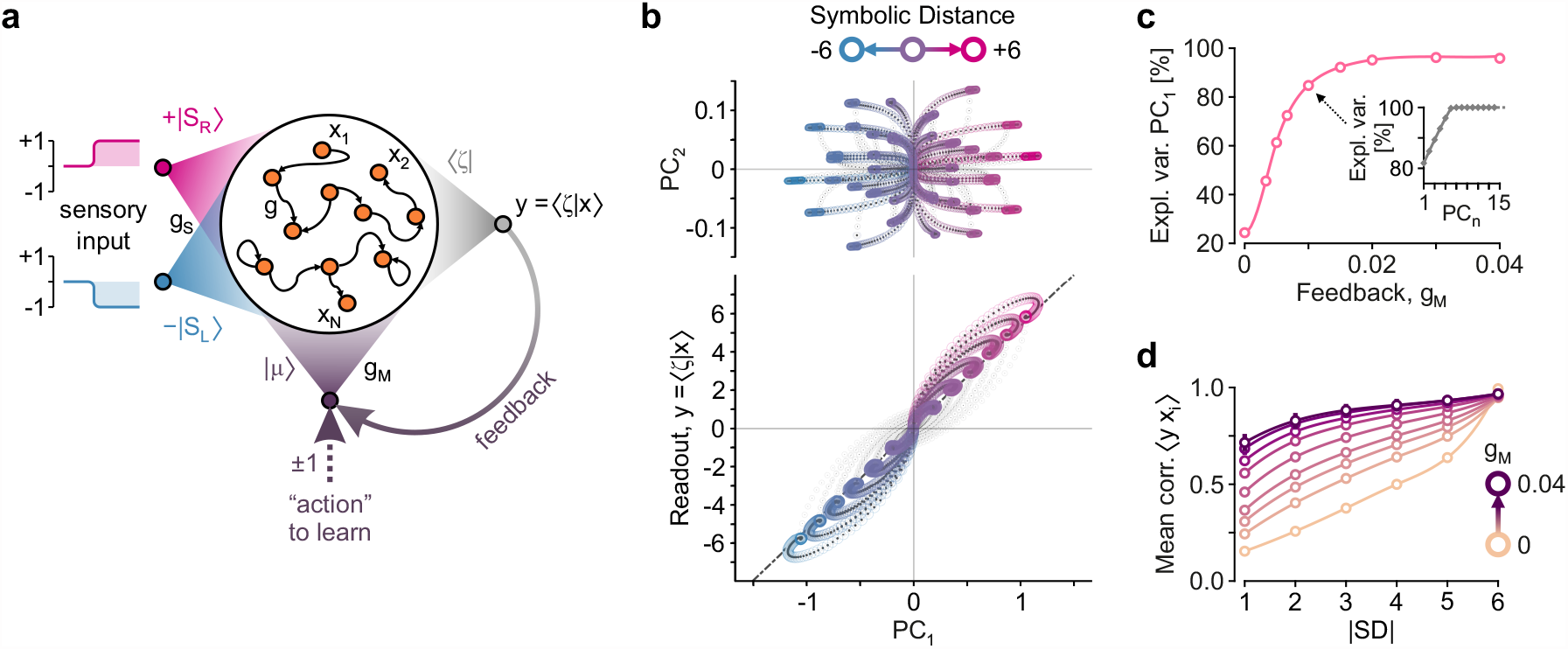
RNN with feedback learn the GML reducing the dimensionality of the latent space. **a**. RNN as in Fig. 4 with the additional input mediated by the weights |*μ*⟩ with strength *g*_*M*_ eventually modulated by the feedback (solid arrow) provided by the readout *y*. During the learning phase only pairs of items with unitary symbolic distance are presented (as in Fig. 3a). The correct action to reach the item with highest rank to the right (left) is represented by the input |+*μ*⟩ (|−*μ*⟩) (dotted arrow). **b**. Neural trajectories of the RNN during the test phase (as in Fig. 3a). The learned GML ⟨*ζ*| results from the pseudo-inverse of the neural activity during the learning phase. Principal component analysis (PCA) is performed on the neural activity of the whole test phase. Trajectories are shown in the planar subspace determined by ⟨*ζ*| and the first PC (bottom), and in the plane of the first and second PCs (top). Colors code for the symbolic distance between the items of the presented pair {*S*_L_, *S*_R_}. Here, the RNN has *N* = 100, |*x*(0)⟩ = 0, *g* = 0.1, *g*_S_ = 0.01 and *g*_M_ = 0.02 (see Methods for other parameters and task details). **c**. Variance explained by the first principal component PC_1_, varying the feedback strength *g*_M_ for a RNN with same parameters as in (b). Inset, variance explained by first 15 PCs for the RNN with *g*_M_ = 0.01. **d**. Mean correlation 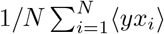 averaged across trials with pairs having the same absolute symbolic distance: |SD| = |*r*_R_ − *r*_L_|. Mean correlations *versus* |SD| are shown for networks with varying feedback strength *g*_M_. Other parameters: *g* = 0.05 and *g*_S_ = 0.01.

In simulation we tested the response of the RNN with feedback to all possible item pairs, finding that it successfully solved the task. Indeed, by performing a principal component analysis (PCA) on the network states, the neural trajectories in the latent plane (PC_1_, *y*) clearly cluster according to the symbolic distance of the presented pairs (Fig. 7b-bottom). Besides, the asymptotic value of the readout lim_*t*→∞_ *y*(*t*) = *y*_∞_ returns the symbolic distance SD = *r*_*k*_ − *r*_*j*_.

For our linear RNN, this can be analytically proven as the asymptotic activity is 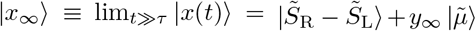. Here 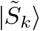 and 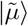 are the inner representations of the presented items and of the motor plan, respectively, transformed by the activity reverberation in the network (see Methods). As a result the GML has to be

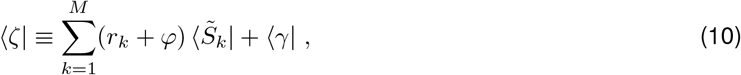

holding for any real *φ* and row vector ⟨*γ*| orthogonal to the subspace where the item representations 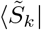 and motor plan 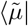 live. This expression is equivalent to the GML derived in Eq. (4), confirming it is a general solution holding also for RNN with feedback. Note that the found ⟨*ζ*| does not depend on the motor plan. Thus, although both the sensory and the motor engrams coexist in the network, they do not interfere.

In Fig. 7b-top it is interesting to note that a richer structure of neural trajectories emerges in the plane of the first two principal components (PC_1_, PC_2_). Trials with same symbolic distance SD now split, unfolding the information about the items in the presented pairs. This is a direct consequence of the fact that representations of item pairs occupy an *M* -dimensional subspace (Fig. 1d), and this information persists in the asymptotic state of the network in Eq. (16). Recurrent activity transforms these internal representations aligning the network state along the axis 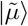, which in turn is the average |*x*_∞_⟩ across trials. For this reason, stronger is the feedback *g*_M_, larger is the variance explained by PC_1_ (Fig. 7c). Not only, as in the above expression for |*x*_∞_⟩ (i.e., Eq. (16) in Methods) the contribution of the motor component is also proportional to the symbolic distance SD = *y*_∞_, also the activity level of the network units are expected to grow according SD. Such correlation is apparent in Fig. 7b-bottom for the example network, and we found it to be weakened by the increase of the feedback strength *g*_M_ (Fig. 7d). This is due to the sigmoidal activation Φ(*h*) limiting the activity *x*_*i*_ to be in the range [− 1, +1].

To summarize, networks with feedback are capable to encode simultaneously both motor-related activity and the same geometric mental line found in more simplistic conditions. The dimensionality of the latent state-space of the network is compressed according to the strength *g*_M_ of the learned feedback. This can limit the degree of correlation between the activity of the network units and the symbolic distance of presented pairs. Remarkably, a similar SD-dependent modulation of the motor-selective activity was found in PMC neurons of monkeys performing a TI task [39].

## 3 Discussion

The brain encodes task relevant information into complex dynamical patterns of neuronal activity, requiring a high-dimensional state space to be folded [9, 44]. These high-dimensional representations have the great advantage to be linearly separable, as random projections of correlated representations tend to be orthogonal in the target high-dimensional neural space, according to the ‘efficient coding hypothesis’ [45, 46]. Linearly separable representations can be indeed categorized *via* relatively simple learning rules eventually leading to a dimensional reduction of the output-potent space [47]. Remarkably, such low-dimensional latent spaces where correlated neural activity is constrained to wander, are pervasively found in associative cortices of many species suggesting it as successful computational strategy preserved across evolution [48, 49, 8].

Here we widen the realm of cognitive functions exploiting such “expansion-compression” computational paradigm to the representations of stimuli and behavioural output in neuronal networks. We indeed prove that the task of arbitrarily ranking a set of *M* abstract items can be always performed looking at the projections of their inner representations onto a suited low-dimensional subspace. Centroids of these projections can be arbitrarily mapped on any monotonic function of the ranks assigned to the items. The subspace where item ranks are encoded works as a “geometrical mental line” (GML), given by a suited linear combination of the item representations. In a network composed by *N* neurons, the GML is not unique as all the *N* − *M* dimensions of the residual state space is unconstrained, leading to an infinite set of solutions. As a result, other network resources can in principle be used simultaneously to encode additional information and/or to perform complementary computation. The existence of a GML thus implies that the dimension of the representational space reduce from *M* to one as a byproduct of the nonlinear dynamics naturally occurring into a recurrent neuronal network, compressing the information about the ranks into a scalar magnitude. The resulting one-dimensional manifold straightforwardly inform about the motor decision (output) to take, which in turn we have shown to be modulated by task difficulty (i.e., different symbolic distances).

Thanks to the fact that our model is based on the linear combination of representations, the GML is highly flexible [9, 50, 51] and different directions can be read out for different arbitrary orders of the same set of items (Figure 2). This supports the idea that in serial reasoning two different representations must be learned [3]: the item representations, which are stored in a suited neural subspaces, and the ordinal information, which depends on the readouts shaped by learning constrained by the task to perform.

Intriguingly, the geometrical mental line successfully faces task sets never seen during the learning phase. This capability to generalize is a hallmark of serial reasoning needed to solve transitive inference task. The formulation of a geometrical solution of a task is the backbone of many cognitive models of serial reasoning [10, 51, 52, 53]. The idea is that in cognitive processes, it is important to consider both the content and the geometrical structure of the neural activity, which fixes the relationships between the relevant variables of the task. In the specific case of transitive inference, the geometric framework underlying the task has widely believed to be a linear workspace [12, 54, 55, 56, 57]. The GML we derived represents this one-dimensional manifold allowing to solve the TI task. Taking into account the intrinsic uncertainty of item representations (sensory noise) and the intrinsic stochasticity of the neuronal activity (thermal noise), simulated recurrent neural networks (RNN) reproduced all the behavioral effects observed in experiments: i) the symbolic distance effect, both in performances and reaction times, and ii) the serial position effect. These results can be explained in terms of a varying signal-to-noise ratio for different couples of items.

Further investigating the same RNN, some predictions arose about different task settings, *e*.*g*. changing the number of training trials or changing the number *M* of items in the learned sequence. This offers an interesting perspective to understand the predictive power of our theoretical framework looking at direct comparisons with behavioral data from experiments [58]. Focusing on the specific case of item sequences with increasing lengths, we expect to see a change in the concavity of the accuracy in the SDE (Fig. 6b) together with a flattening of the performances for central items in the SPE. A result which could inform about the limitations of the network in the capacity to store serial-order information. A limitation which in experiments is often worked around resorting to the so-called ‘linking chain paradigm’ [36, 59]).

Furthermore, we proved that the GML can be naturally embedded into a RNN designed to make motor decisions. In this case a one-dimensional manifold naturally emerges in the latent space visited by network activity, which is aligned motor plan encoding the decision to take. Such dynamical organization of the network dynamics accounts for the expected modulation of single-unit activity observed in the premotor cortex (PMC) of monkeys, according to symbolic distances between presented items [39]: stronger is the response, larger is the number of units in the RNN exceeding a given activity threshold.

Starting from this evidence we speculate that PMC might be an ideal candidate to implement the GML solution when the transitive inference task is learned and successfully performed. We can also speculate that this mental line representation might play a role well before movement onset, as it has been shown that in the delay period following the presentation of the item pairs, the decision is already taken and the symbolic distance is no more relevant [60].

Notably, we also found that in addition to the symbolic distance, the GML allows our network models to provide as a read out the joint rank (i.e., the rank sum) of item pairs. This happens if the item representations are summed (|*S*_R_ + *S*_L_⟩) instead of being subtracted (|*S*_R_ − *S*_L_⟩). Interestingly, in the posterior parietal cortex of non-human primates neuronal activity was found to correlate with the joint rank despite the fact that it was not a relevant information to solve the TI task [61]. According to this evidence, a reasonable hypothesis to test is that different sensory pathways might coexist implementing the sum and the difference of item representations presented in pair. In this way cortical networks would be spontaneously ready to respond according to both the symbolic distance and the joint rank, a kind of primitive algebraic competence.

Regarding the linear combination of item representations determining the GML ⟨*ζ*| in Eq. (3), it is important to remark that the coefficients (i.e., the projection values) of such combination are given by the item ranks *r*_*k*_. This is due to fact that the TI task is learned on trials with item pairs with |SD| = 1. However, depending on the training procedure adopted in the serial learning task, the response can be shaped as a function *f* (*r*_*k*_) of the item ranks, as emphasized in Eq. (3). This function can be also modulate by the specific amount of reward the subject receives for different item pairs. In the experimental literature there are several possible modifications of the reward schedule or in the training procedure of specific pairs of items that can change the projection values embedded into the GML, without affecting the capability to encode the correct order in the sequence ([38, 27, 62]). Alternatively, also the number of trials composing the training phase can have a similar impact, eventually leading to different slopes in the correlation shown in Fig. 3e. We can also imagine a task setting leading to the logarithmic function *f* (*r*_*k*_) = log(*r*_*k*_), as in the case of an over-training of the low-rank items as hypothesized in the ordering of numerical quantities [31]. In these conditions, the projection values of ⟨*ζ*| are no more equally spaced: the first items have a greater distance along the GML. This might explain some behavioral effects, such as those following the Weber’s principle formalized as Fechner’s law [63, 64], giving rise to the well-known primacy effect possibly observed in various serial learning tasks (e.g., in the simultaneous chain task [65, 15]).

Finally we remark that several models have been proposed to explain how a mental scheme of the items rank could be learned by humans and animals [30, 31], and what is the role of the reward in this process [66, 58]. An intriguing and open question is then, whether such theoretical frameworks together with the one we presented here, are capable to predict effects not yet observed sharpening their range of applicability, and thus shading further light on the brain mechanisms underlying serial thinking.

## Methods

### RNN dynamics and its linearization

Recurrent neural networks (RNN) are composed of *N* units (*N* = 100, unless otherwise specified in the main text). The *j*-th unit has activity state *x*_*j*_(*t*) = {|*x*(*t*)⟩}_*j*_ evolving according to the first-order dynamics

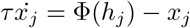

for any *j* ∈ [1, *N*] ⊂ℤ The decay time constant *τ* = 0.1s and the activation function Φ(*h*_*j*_) ≡ tanh(*h*_*j*_) is the same for all units. The synaptic input *h*_*j*_ = {|*h*⟩}_*j*_ is the weighted sum

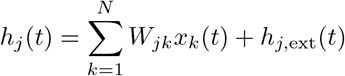

where *W*_*jk*_ are the element of the synaptic matrix **W** ∈ ℝ^*N*×*N*^ sampled randomly from a Gaussian distribution with 0 mean and standard deviation 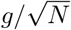 is the external input received by the unit *j*. The network dynamics is integrated relying on the first-order Euler-Maruyama method with time step *dt* = 0.1*τ* to take into account the possible sources of noise in the input detailed in the main text and in the following. In matrix form the above dynamics can be compactly written as

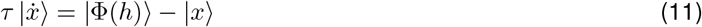

With

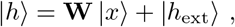

where the ‘kets’ |·⟩ ∈ ℝ^*N* ×1^ are column vectors.

During the serial ranking tasks discussed in the main text, the presentation of items on the screen is modeled as a modulation of the external input. More specifically, the single *k*-th item elicits a sensory input |*h*_ext_⟩ = *g*_S_ |*S*_*k*_⟩ with strength *g*_S_ set to 1 when not specified otherwise. The vector elements *S*_*ki*_ = {|*S*_*k*_⟩}_*i*_ are i.i.d. random variables with zero mean and variance 1*/N*. The ‘bras’ ⟨*S*_*k*_| ∈ ℝ^1×*N*^ are the row vectors transposed of |*S*_*k*_⟩, such that in the large-network limit, the inner product is

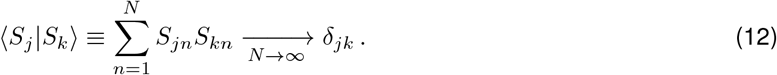

When two items are simultaneously presented as in the TI task, the input received by the network is a linear combination of the isolated stimuli: |*h*_ext_⟩ = *g*_S_ (|*S*_*j*_⟩ − |*S*_*k*_⟩) ≡ *g*_S_ |*S*_*j*_ − *S*_*k*_⟩. For the sake of simplicity, the item *k* presented on the left side of the screen has negative strength, while the *j*-th located on the right contributes as it would be alone. Such peculiar modeling choice is not critical and can be generalized as shown below (Subsection ‘GML with arbitrary item representations’).

When both the spectral radius *g* of **W** and the strength *g*_S_ of the sensory input are small (*g, g*_S_ ≪ 1), the dynamics of the RNN can be linearized. Indeed, under these conditions assuming a sensory and thermal noise of the same order of magnitude (see below), the synaptic input *h*_*j*_ is expected to be of the same order. Hence, the activation function can be approximated as Φ(*h*_*j*_) ≃ Φ(0) + *h*_*j*_Φ′(0) = *h*_*j*_, and Eq. (11) reduces to

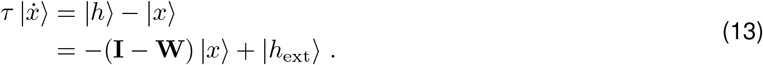

As above, at rest |*h*_ext_⟩ = |0⟩, while when the pair of items *j* and *k* appears to the right and to the left of the screen, respectively, |*h*_ext_ = *g*_S_ |*S*_R_ − *S*_L_⟩ with *S*_R_ = *S*_*j*_ and *S*_L_ = *S*_*k*_. All the RNN studied in this work are linear and as such follow the dynamics (13).

### Linear RNN and motor decision

In the linear RNN introduced in Sect. 2.4, we impose that the response unfolds along an independent 1-dimensional manifold encoding the motor plan. To this purpose an additional motor-related input *g*_M_ |*μ*⟩ is added to |*h*_ext_⟩ further modulated by the expected response *y*′(*t*) leading to the linear dynamics

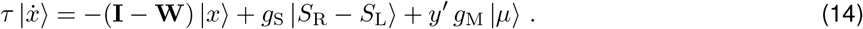

Here *g*_S_ and *g*_M_ modulate the strength of the sensory and motor-related inputs, respectively. As the item representations |*S*_*k*_⟩, the ‘motor’ axis |*μ*⟩ is extracted randomly making all these engrams orthogonal (i.e., ⟨*S*_*k*_|*μ*⟩ = 0 for any item *k*) in the limit of infinitely large networks (*N* → ∞).

In the learning phase involving only the set of pairs {*S*_*j*_, *S*_*k*_} with unitary symbolic distance, i.e., |SD| = |*r*_*j*_ − *r*_*k*_| = 1 (see Fig. 2a), the expected response is *y*′ = +1 (−1) when the item to reach is on the right (left). As in the standard reservoir computing framework [67], if the GML ⟨*ζ*| solving the task *y*′ = ⟨*ζ* | *x*⟩ exists, it results from a linear regression of the activity leading to a new synaptic matrix

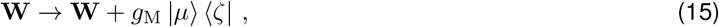

where the rank-1 matrix *g*_M_ |*μ*⟩ ⟨*ζ*| represents the changes induced by learning.

In simulations, we computed the mental line ⟨*ζ*| during the learning phase, resorting to the pseudo-inverse of the neural activity of the RNN without feedback (i.e., without re-injecting ⟨*ζ* | *x*⟩ as factor of |*μ*⟩). We then updated the synaptic matrix according to Eq. (15), eventually testing the response of the RNN with feedback to all possible item pairs. In this testing phase only the sensory input *g*_S_ |*S*_R_ − *S*_L_⟩ is provided, as the motor plan spontaneously emerged from the recurrent activity of the linear RNN.

From Eq. (14) the asymptotic state of the network following the presentation of a pair of items can be carried out by imposing 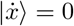, which results to be

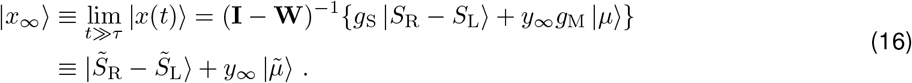

Here 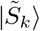 and 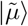 are the inner representations of the presented items and of the motor plan, respectively, transformed by the activity reverberation in the network.

Even in this general case it is easy to prove that the geometric mental line ⟨*ζ*| solving the TI task is given by a linear combination of row vectors 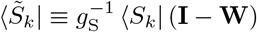. Indeed, for such vectors the following relationships hold: 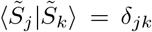 and 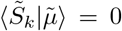. With that, by imposing ⟨*ζ*|*x*_∞_⟩ = *r*_R_ − *r*_L_ for all the item pairs with |SD| = 1, an infinite set of geometric mental lines exist and they are

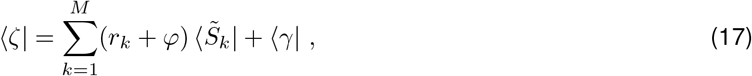

i.e., Eq. (10) reported in the main text. Note that by setting **W** = 0 and *g*_S_ = 1 the simple linear-Perceptron configuration introduced in the main text is recovered, eventually making the above expression equivalent to Eq. (3).

From Eq. (14) we can work out the evolution in time of the readout *y*(*t*) and the projection *z*(*t*) ≡ ⟨*μ*|*x*⟩ expected to faithfully represent the loading of the PC_1_. Indeed applying to both hand sides of such equation the ‘bra’ ⟨*μ*| and ⟨*ζ*| we obtain the following analytical approximations

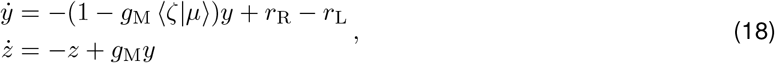

where we assumed that both ⟨*ζ* |**W**|*x*⟩ and ⟨*μ* |**W***x*⟩ are approximately equal to 0 as we verified numerically in simulation. From this expression it can be noted that the network activity starting from |*x*⟩ = 0 initially unfolds being driven exclusively by the information |*S*_*k*_ − *S*_*j*_⟩. Afterward, the contribution from the activity *z*(*t*) along |*μ*⟩ starts to play a role being modulated by a *y*(*t*), as it is apparent from Fig. 7b-bottom looking at the slopes of the trajectories around PC_1_ = 0. We finally remark that the asymptotic value of *z*_∞_ = lim_*z*→∞_ *z*(*t*) returns to be fully correlated with the projection of the network activity onto the GML: *z*_∞_ = *g*_M_*y*_∞_ = *g*_M_(*r*_R_ − *r*_L_).

### Linear RNN with sensory and thermal noise

In Sect. 2.4, the dynamics (13)of the above linear RNN incorporates units receiving a stochastic input. This is done by adding to the input |*h*⟩ a ‘thermal’ noise |*ξ*^(T)^⟩ uncorrelated in time whose elements 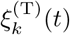 are i.i.d. random variables extracted from a Gaussian distribution with zero mean and an arbitrary variance 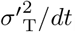. Sensory input in this network configuration is given by the superposition of the contributions due to the presented items (|*h*_ext_⟩ = *g*_S_ |*S*_R_ − *S*_L_⟩) eventually leading to the linearized dynamics

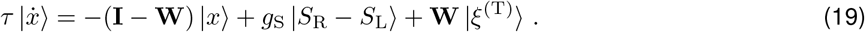

In this framework, the readout weights ⟨*ζ*| given by Eq. (3) make the network capable to solve on average the TI task provided that they are rescaled by 1*/g*_S_ (i.e, 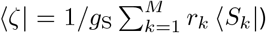. Indeed, applying to both hand sides of Eq. (19) the rescaled GML we obtain the stochastic dynamics

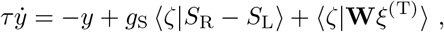

where we neglected the *𝒪*(*g*) terms approximating **I** − **W** ≃ **I**. Here ⟨*ζ*|**W***ξ*^(T)^⟩ results to be a Gaussian random variable with zero mean and variance 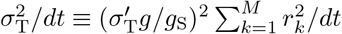 which for *r*_*k*_ = *k* gives

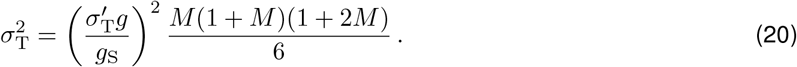

This because the vector and matrix elements *S*_*ki*_, *W*_*ij*_ and 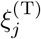 are i.i.d. random Gaussian variables with zero mean and variance 1*/N, g*^2^*/N* and 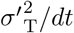, respectively. Considering that for the chosen readout weights *g*_S_ ⟨*ζ*|*S*_R_ − *S*_L_⟩ = *r*_R_ − *r*_L_, the stochastic dynamics of the readout *y*(*t*) eventually reduces to Eq. (7) in Sect. 7:

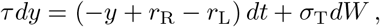

where *y*(*t*) is driven by a Gaussian white noise *dW* (*t*) with zero mean and covariance 𝔼[*dW* (*t*)*dW* (*t*′)] = *δ*(*t* − *t*′)*dt*.

In addition to thermal noise, the model incorporates sensory noise to account for the variability in encoding visual stimuli across trials. The actual contribution of the presented pair of items in a generic trial *n* is given by: 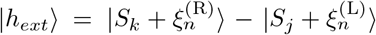, where 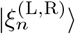 is a vector of i.i.d. Gaussian values with zero mean and standard deviation 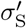. This leads to a random contribution to the item direction |*S*_*k*_⟩ with a variance of 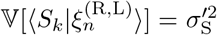, and for the projection along the GML 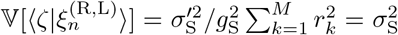. Given *r*_k_ = *k*, the following relationship holds

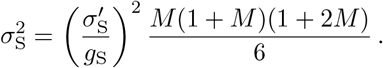

In order to preserve the linear approximation of the dynamics, the effective intensity of sensory contribution 𝕍[*g*_S_ |*h*_*ext*_⟩] is fixed rescaling *g*_S_ by a factor 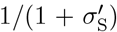. Finally the stochastic dynamics of the readout *y*(*t*), considering the two noise sources, is updated as follows

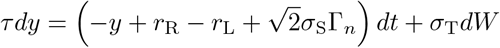

where Γ_*n*_ is an independent random Gaussian variable with zero mean and unit variance.

### Linear RNN parameters and task design

Regarding the simulated linear RNN, they have random connectivity with spectral radius *g* = 0.05 and strength of sensory input *g*_S_ = 0.2*g*. Each item is schematized by injecting a square-wave current lasting 1.5*s*, smoothed by a Gaussian-weighted moving average filter and followed by a relaxation time of 3*s*. The state history associated with the items pairs with adjacent rank (|SD| = 1) randomly presented 30 times each, is used to estimate the vector of readout weights through the pseudo-inverse approach, eventually determining a motor response the activity of the readout unit *y* crosses a threshold value of 0.5. The accuracy of the responses is tested across the whole set of possible item pairs (i.e., the training set) presented for 20 times.

The task visual stimuli are simulated by stimulating groups of nodes with a square wave current, positive when the stimulus is placed on the right side of the screen, negative when it is placed on the left. As previously shown, two different sources of noise are considered: sensory 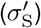 and thermal noise 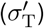. The interplay between the two sources in the operation of the recurrent neural network is illustrated in Fig. 5.

The network is trained on couples of nearest-neighbors symbols, computing the linear readout such that the state of the system gives +1 when the target item is on the right and −1 when it is on the left. The resulting output is an accumulation process that triggers a choice once a threshold of 0.5 is overcome. If the signal does not reach the threshold within the 1.5 s of the square wave stimulation the trial is not taken into account in the performances computation.

### GML with arbitrary item representations

For the sake of simplicity in the main text we assumed that item representations on one side of the screen (left or right) are associated to opposite network states, such that ⟨*S*_*k*L_ |*S*_*k*R_⟩ = −1 for any *k* ∈ [1, *M*]. Here we prove that even when this assumption is relaxed, the GML is still a linear combination of the decoding weights discriminating both the presence and the position of the *k*-th item on the screen, as in Eq. (3). To this purpose let’s consider the general case of independent item representations (⟨*S*_*kα*_|*S*_*jβ*_⟩ = 0 for any k≠ *j* and *α, β* ∈ {L, R}). This choice is justified by the efficient coding hypothesis such that sensory information is orthogonalized via random projects onto a high-dimensional representational space [45, 46]. However, it is reasonable to expect that some degree *ρ*_*k*_ of correlation exists between the representations of the same item appearing in different positions of the screen: ⟨*S*_*k*,L_|*S*_*k*,R_⟩ = *ρ*_*k*_. In this case the GML solving the TI task from the learning phase must solve the following set of 2(*M* − 1) equations

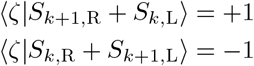

for any *k* ∈ [1, *M* − 1].

Making also in this case the ansatz 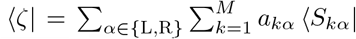 work out the coefficients, the above set of equations allows to work out the coefficients *a*_*kα*_ for the decision to choose the highest item on the right

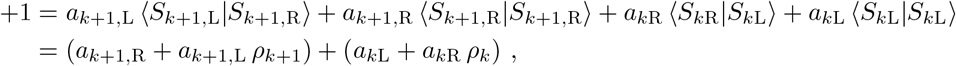

or on the left

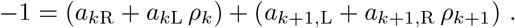

Summing these two equations we obtain

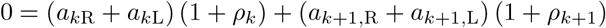

which are always solved if *a*_*k*L_ = − *a*_*k*R_ for any *k* ∈ [1, *M*]. This together with the difference of the previous two equations leads to have

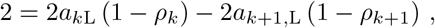

eventually allowing us to write the solution

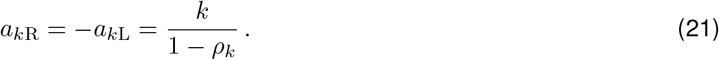

Given these coefficients, the generalized GML result to be

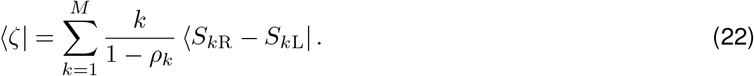

Note that in the simplified case discussed in the main text (*ρ*_*k*_ = −1) the GML Eq. (3) is recovered as ⟨*S*_*k*L_| = − ⟨*S*_*k*R_|.

## Dimensionality analysis

Fig. 7 depicts the outcomes of the principal component analysis (PCA) conducted on the network activity simulated during the testing phase of the task. To perform PCA, we first centered the sampling matrix *T* × *N* at zero by subtracting its time averages. Here *T* denotes the number of time steps of the test phase and *N* signifies the number of nodes in the network. Then, we computed the associated covariance matrix and its eigenvectors, which represented the principal components, and sorted them based on the magnitude of the corresponding eigenvalues. Notably, the projection of the network state onto the eigenvector linked with the maximum eigenvalue equates to the projection onto the first principal component.

## Distributions analysis

The distributed representation of items onto the GML, as shown in Fig. 5b, has been estimated by presenting to the network by a single item per trial for 30 trials each.

## Code availability

All analyses were implemented using custom Matlab code. Codes regarding key findings will be shared upon specific request.

## Acknowledgments

We are deeply indebted to E. Brunamonti and S. Ferraina for having introduced us to the transitive inference task and for the many stimulating discussions. Work partially funded by EU H2020 Research and Innovation Programme, Grant 945539 (HBP SGA3) to MM. Preliminary results of this work have been presented in [68, 69].

